# The ITPRIPL1- CD3ε axis: a novel immune checkpoint controlling T cells activation

**DOI:** 10.1101/2022.02.25.481189

**Authors:** Shouyan Deng, Yiting Wang, Jean-Philippe Brosseau, Yungang Wang, Jie Xu

## Abstract

The immune system is critical to fighting infections and disease. The molecular recognition of harmful entities takes place when antigen-presenting cells (APC) harboring major histocompatibility complex (MHC) molecules bound to peptides derived from harmful antigens (ligand) dock on specific T cell receptor (TCR)-CD3 complex (receptor) at the surface of CD8+ T cells. The discovery of a general immune checkpoint mechanism to avoid the harmful impact of T cell hyperactivation provoked a paradigm shift. The clinical relevance of this mechanism is highlighted by the fact that PD-1 and PD-L1 inhibitors are very effective at boosting immune reactions. Still, immune evasion frequently happens. The observation that some PD-1/PD-L1 negative tumors have a poor immune response opens the door to identifying a novel immune checkpoint mechanism. Here, we discovered that ITPRIPL1, a gene with unknown function, impairs T cell activation. Surprisingly, we found that CD3ε is the direct receptor of ITPRIPL1. This novel immune checkpoint was validated as a drug target using ITPRIPL1 KO mice and monoclonal antibodies. Thus, targeting the ITPRIPL1-CD3e axis, especially in PD-1 - PDL-1 negative patients, is a promising therapeutic strategy to reduce immune evasion.

## Introduction

The immune system is key to fighting infections and disease. Several white cells, including macrophages, neutrophils, and lymphocyte T (T cells), ultimately kill the defective cells and/or foreign material [1]. In some pathologies such as cancer, the immune system is not sufficient to overcome the defective cells [2]. Reprogramming (boosting) the immune system to promote disease remission has been the holy grail of many immunologists for the last few decades [3, 4].

The molecular recognition of harmful entities takes place when antigen-presenting cells (APC) harboring major histocompatibility complex (MHC) molecules bound to peptide derived from harmful antigens (ligand) dock on specific T cell receptor (TCR)-CD3 complex (receptor) at the surface of CD8+ T cells [5, 6]. The discovery of a general immune checkpoint mechanism to avoid the harmful impact of T cell hyperactivation provoked a paradigm shift [7]. One of the most studied immune checkpoints is the PD-L1 (ligand on APCs)/ PD-1 (receptor on T cells) checkpoint [8]. One way to escape the immune system is to stimulate this checkpoint mechanism [9]. Conversely, the clinical relevance of this mechanism is highlighted by the fact that PD-1 and PD-L1 inhibitors are very effective at boosting immune reactions [10]. However, the observation that many “hot” tumors are negative for PD-L1 opens the door to identifying a novel immune checkpoint mechanism [11-13]. In this sense, many additional immune checkpoint receptors and ligands have been validated as drug targets [14-18]. Still, immune evasion frequently happens [19, 20]. In the classic two-signal model of T-cell activation, the primary signal (signal one) is triggered by the ligation of the peptide-MHC to the TCR-CD3 complex, where the TCR represents the only receptor involved [21, 22]. Signal two is generated by the transduction of co-activators or co-inhibitors in the immune synapse following a calcium influx [23, 24]. Our quest to identify a novel immune checkpoint leads us to the discovery and drug target validation. ITPRIPL1 is a gene with previously uncharacterized functions, and CD3ε is surprisingly working as a direct receptor of ITPRIPL1 to impair T cell activation.

## Results

### *ITPRIPL1* is associated with immune evasion

Functionally redundant genes frequently display a mutually exclusive pattern of molecular alterations in disease. In the context of immune evasion, several immune checkpoints are functionally redundant to PD-L1/PD-1[25]. To uncover novel immune evasion mechanisms, we decided to mine the transcriptome of thousands of cancer cell lines to identify transmembrane protein-encoding genes that show significant mutual exclusive pattern with CD274 (PD-L1) expression. Doing so, we identified *ITPRIPL1* (Fig.1A and supplementary Fig.1A), a gene with unknown functions.

**Fig. 1.**
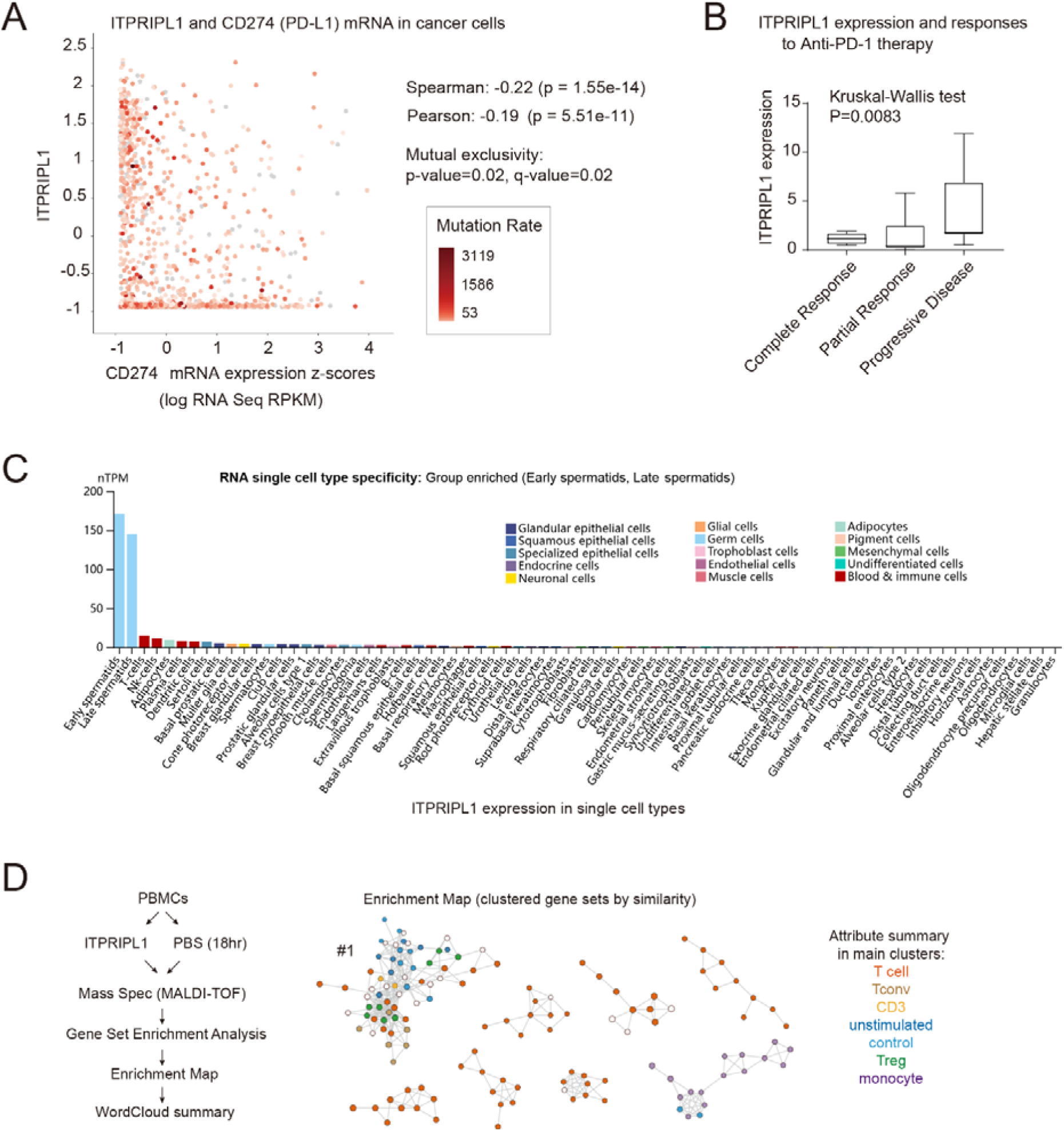
*ITPRIPL1* is associated with immune evasion. **A**, Scatter plot illustrating the mutual exclusivity of ITPRIPL1 and PD-L1 as judged by analyzing the transcriptomic datasets from the Cancer Cell Line Encyclopedia (CCLE) (FDR-adjusted P<0.05). **B**, Bar graph of the ITPRIPL1 mRNA expression showing a negative correlation between ITPRIPL1 expression and anti-PD-1 therapeutic effects (FDR-adjusted P<0.05) after re-analysis of the GSE78220 dataset. **C**, Bar graph of ITPRIPL1 expression in different cell types. Single cell expression atlas showing physiological ITPRIPL1 expression is associated with immunogenic tissues. **D**, Mass spectrum enrichment map analysis suggests a role in suppressing T cells signaling for ITPRIPL1.

If it is true that *ITPRIPL1* is functionally redundant to the PD-L1-/PD-1 axis, then ITPRIPL1 should take over as an immune checkpoint where *PD-L1*/*PD-1* does not. In this sense, we examine a transcriptomic dataset of tumors derived from cancer patients who displayed a varying degree of response to anti-PD-1 therapy. As predicted, *the ITPRIPL1* expression level was significantly higher in the PD-1 insensitive group (Fig.1B).

If it is true that ITPRIPL1 contributes to tumor immune evasion, then under physiological conditions *ITPRIPL1* should be expressed in immune privileged organs. This is based on the fact that PD-L1 displays the highest expression level in placenta and plays a key role in maternal-fetal immune tolerance [26]. Indeed, ITPRIPL1 is enriched in spermatid in the testis, the strongest immune privileged organ [27] as judged by the Single Cell Expression Atlas (Fig.1C). Overall, ITPRIPL1 is a molecule specifically expressed in immune privileged tissues, “hot” tumors without PD-L1 expression, and non-responding tumors to PD-1 blockade therapy.

To validate whether ITPRIPL1 affected the immune system, we first evaluate the capacity of exogenous ITPRIPL1 purified protein to modify T cell signaling in human peripheral blood mononuclear cells (PBMCs), a widely used source of immune cells. Proteome-wide profiling of ITPRIPL1-treated PBMCs followed by Gene Set Enrichment Analysis on differentially expressed proteins pointed to suppressive signaling in T cells (Fig.1D, supplementary Fig.1C and Table.1).

### *ITPRIPL1* impairs T cell function *in vitro* and *in vivo*

To further determine whether ITPRIPL1 impairs T cell cytotoxicity, we performed a series of T-cell killing assay using HCT-116 cells as the killing target of T cells. T cells effectively increase apoptosis of HCT-116 whereas HCT-116 stably overexpressing ITPRIPL1 (HCT116-ITPRIPL1 OE cells), and HCT-116 knockout for ITPRIPL1 (HCT116-ITPRIPL1 KO cells) decrease and increase, respectively, the apoptotic rate (Fig.2A, Supplementary Fig.2A).

**Fig. 2.**
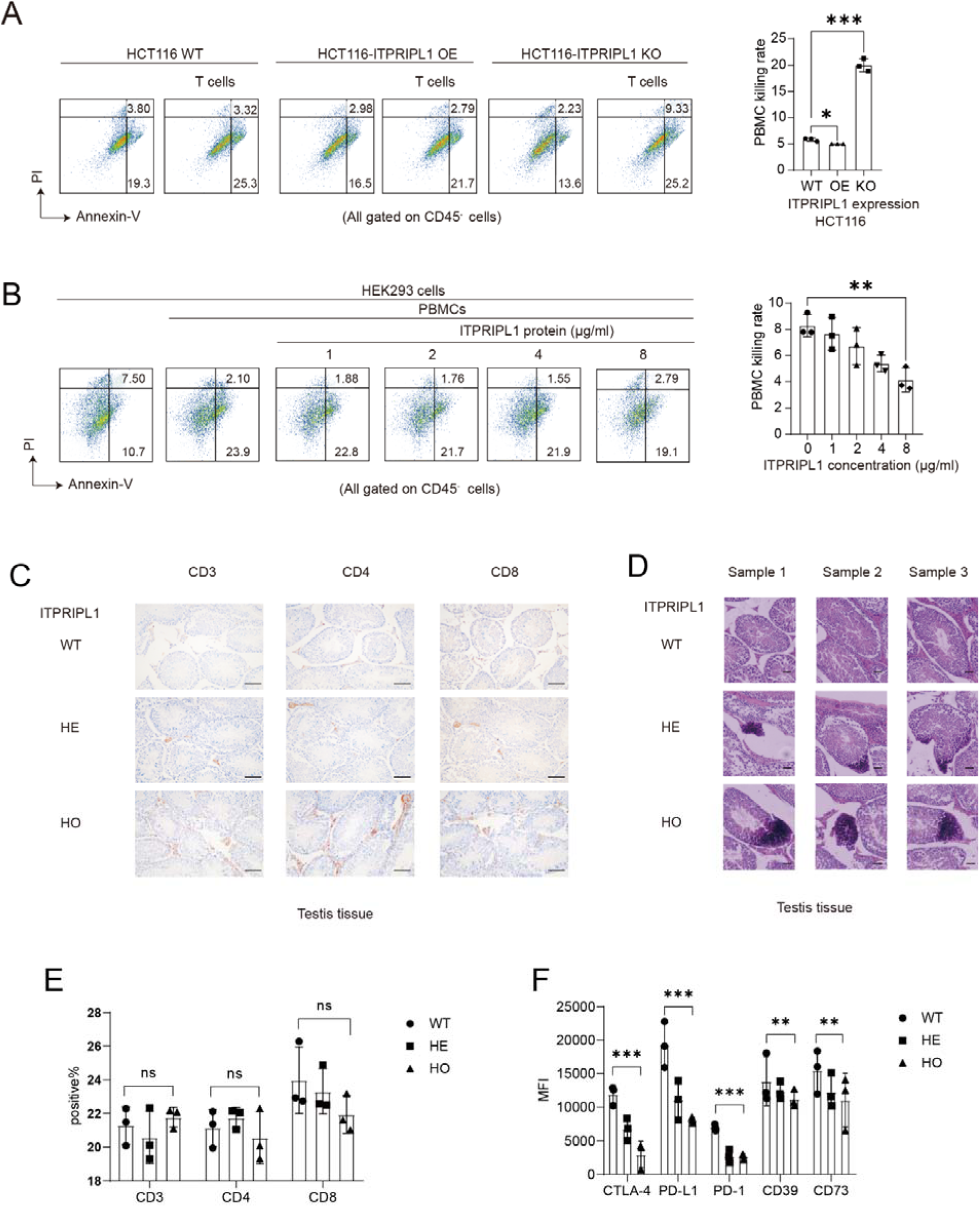
ITPRIPL1 impairs T cell function *in vitro* and *in vivo*. **A-B**, T-cell killing assay showing the immune evasion of (**A**) HCT116 cells and (**B**) HEK293 is dependent upon ITPRIPL1 (**A**) endogenous and (**B**) exogeneous expression level (n=3). **C-D**, Histological evaluation of testis from ITPRIPL1 WT, HE and HO mice at 100X magnification showing (**C**) more CD3^+^/CD4^+^/CD8^+^ cells infiltration after ITPRIPL1 knockout (n=3). and abnormal testis tissue as judged by Hematoxylen and Eosin staining (n=3). **E**, Bar graph illustrating the expression level of CD3, CD4 and CD8 expression in ITPRIPL1 WT, HE and HO mice by FACS. No significant changes in the total was observed (n=3). **F**, Bar graph illustrating the mean fluorescence intensity (MFI) of a panel of immune checkpoint markers by FACS. It shows substantial decrease in inhibitory immune molecules including CTLA-4, PD-L1, PD-1, CD39, CD73 after ITPRIPL1 knockout (n=3). Data are mean ± s.d. **P<0.05, ***P<0.001, ****P<0.0001. Two-tailed Student’s t-test.

To determine if the exogenous addition of ITPRIPL1 was sufficient at blocking the T-cell killing capacity, we performed the T-cell killing assay on HEK293 as a killing target and added an increasing amount of purified ITPRIPL1 protein. The results indicate that the dose-dependent addition of ITPRIPL1 protein is associated with a decrease in HEK293 necrosis/apoptosis rate as judged by the lower annexin-V^+^ signal (Fig.2B). Altogether, it provides strong experimental evidence supporting a negative correlation between ITPRIPL1 expression level in APCs and T cell cytotoxicity.

To further study the *in vivo* impact of ITPRIPL1 protein, we applied CRISPR/Cas9 technique on C57BL/6J mice to build ITPRIPL1-heterozygous knockout (HE) mice. The mice genotype was confirmed by PCR (Supplementary Fig.2B). Next, we bred these HE mice to obtain viable and healthy ITPRIPL1-homozygous knockout (HO) mice. Since spermatids are strongly immunogenic and highly express ITPRIPL1, we first investigate the impact of knocking out ITPRIPL1 in testis immune cells. To do so, we collected testis in a cohort of adult WT, HE and HO mice and performed CD3/CD4/CD8 IHC staining. The IHC positive CD3/CD4/CD8 status inside the testis pointed to upregulated immunoreactions (Fig.2C). Gross examination of the testis of HO mice showed specific abnormalities (Fig.2D), and examination of the sperm from HO mice under the microscope revealed abnormal spermatocyte motility, which may indicate enhanced immune reactions inside the reproductive glands (Supplementary Fig.2C).

To investigate the impact of knocking out ITPRIPL1 on their general immune status, we evaluated the expression level of a panel of classic immune checkpoint markers in PBMCs harvested from WT, HE and HO mice. Similar to the findings in VISTA-deficient mice [28], the total CD3/CD4/CD8 percentage was not changed compared to WT mice (Fig.2E). In sharp contrast, the representative T cell exhaustion markers, including CTLA-4, PD-L1, PD-1, CD39, CD73, were markedly decreased in the HO and HE mice groups (Fig.2F), indicating a rescue from the T cell exhaustion state by knocking out ITPRIPL1. This is reminiscent of the LAG-3 (immune checkpoint) deficient mice [29]. ITPRIPL1 KO did not significantly interfere with the general health status as judged by histopathological evaluation of different organs (heart, liver, spleen, brain, salivary gland, pancreas) (Supplementary Fig.2D). Thus, the *in vivo* studies collectively revealed a negative immunoregulative role mediated by ITPRIPL1 under physiological circumstances, and ITPRIPL1 KO activates T cells but did not significantly influence the general health status of the mice.

### CD3ε as a *bona fide* ITPRIPL1 receptor

ITPRIPL1 expression correlated with immune evasion and impairs T cell activity *in vitro and in vivo* suggesting that ITPRIPL1 is binding to a putative ITPRIPL1 receptor on CD8+ T cells. All TCR-CD3 complex acting as receptors for pMHC-ligand ultimately converge to the activation of the NFκB/NFAT family of transcription factors upon calcium entry following the engagement of TCR. Therefore, we used the Jurkat-NFκB/NFAT (Jurkat-dual system: a reporter assay with luciferase expression under an NFκB/NFAT promoter in T cells [19,20]) to show that ITPRIPL1 is acting directly on T cells through a yet to be identified receptor. Indeed, we observed a decrease in luminescence upon ITPRIPL1 stimulation, indicating that ITPRIPL1 can repress essential signaling downstream of the TCR-CD3 complex. Meanwhile, ITPRIPL1 can substantially downregulate the intracellular calcium signal within Jurkat cells, urging us to investigate into the immune checkpoints associated with calcium influx (Fig.3A). This result encouraged us to identify the putative ITPRIPL1 receptor by screening a series of immune checkpoints. Surprisingly, we discovered that only CD3ε interacted with ITPRIPL1 by ELISA (Fig.3B). We repeat the Jurkat-dual assay with an increasing concentration of purified CD3ε protein to validate this finding. The results showed that the effect of ITPRIPL1 can be counteracted by CD3ε protein and hence, demonstrated that ITPRIPL1 makes direct contact with CD3ε (Fig.3C). To investigate this interaction in detail, we performed an ELISA between ITPRIPL1 and CD3ε by coating different concentrations of ITPRIPL1 to interact with the CD3ε recombinant protein. The sigmoid binding curve indicates that ITPRIPL1 binds to CD3ε in a concentration-dependent manner, with an approximate EC50 of 0.2553μg/ml (Fig.3D). We also measure the binding kinetics assays by Octet. The results of which suggested a relatively weak but affirmative binding between ITPRIPL1 and CD3ε, with KD=3.17e-07(M), ka=6.50e+03(1/Ms), and kd=2.07e-03(1/s) (Fig.3E). Thus, CD3ε is a *bona fide* ITPRIPL1 receptor.

**Fig. 3.**
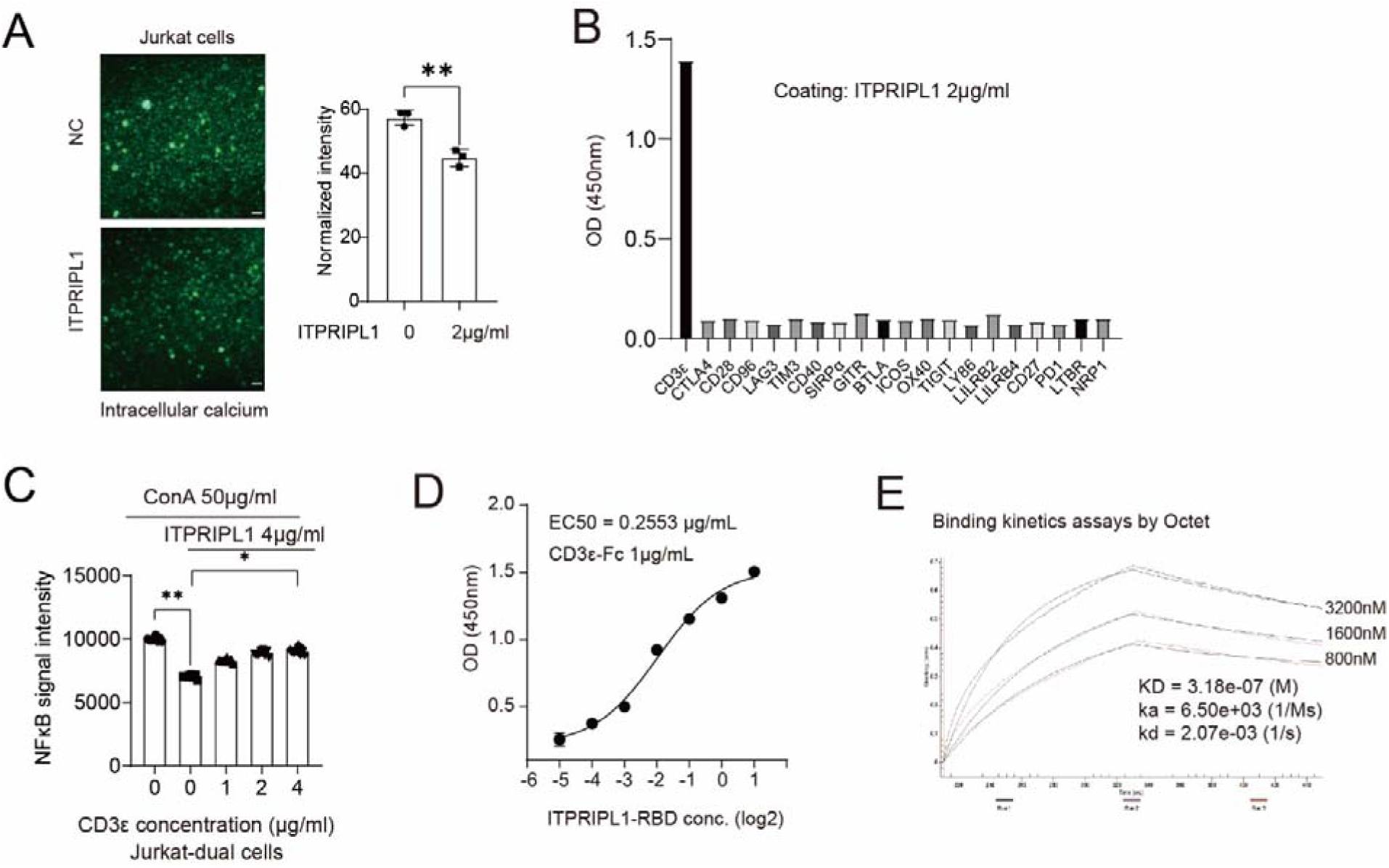
CD3ε as a *bona fide* ITPRIPL1 receptor. **A**, Intracellular calcium analysis in Jurkat cells revealed T cell signal inhibition by ITPRIPL1 at 40X magnification (n=3). **B**, ELISA screening of potential ITPRIPL1 interactors revealedCD3ε. **C**, Jurkat-dual cells luciferase assay revealed the NFκB signal downregulation by ITPRIPL1 was dependent on ITPRIPL1-CD3ε interaction (n=6). **D**, ELISA binding curve of ITPRIPL1-CD3ε (n=2). **E**, Octet showing the association-disassociation curve between ITPRIPL1 and CD3ε with the kinetics constants (n=3). Data are mean ± s.d. **P<0.05, ***P<0.001, ****P<0.0001. Two-tailed Student’s t-test.

### Targeting ITPRIPL1 using a monoclonal antibody de-repress T cells activation

Finally, we developed an ITPRIPL1 monoclonal antibody to restore anti-tumor immunity by blocking the ITPRIPL1-CD3ε interaction. The 13B7A6H3 (13B7) was selected among more than a hundred of clones for its strong binding affinity and specificity (Supplementary Fig.3A-D), and it showed the greatest ability to block ITPRIPL1-CD3ε interaction (Fig.4A). The exact 13B7 peptidic sequence responsible for the binding to ITPRIPL1 was also determined (Supplementary Fig. 3E, Supplementary Table 3). To functionally validate the 13B7 monoclonal antibody, we performed the T-cell killing assay once again. We observe that the *in vitro* T cell cytotoxicity and immune suppression can be reversed by the 13B7 monoclonal antibody (Fig.4B). Thus, we developed a monoclonal antibody specifically targeting ITPRIPL1 and found that this antibody can reverse the *in vitro* function of ITPRIPL1 by blocking the ITPRIPL1-CD3ε interaction.

**Fig. 4.**
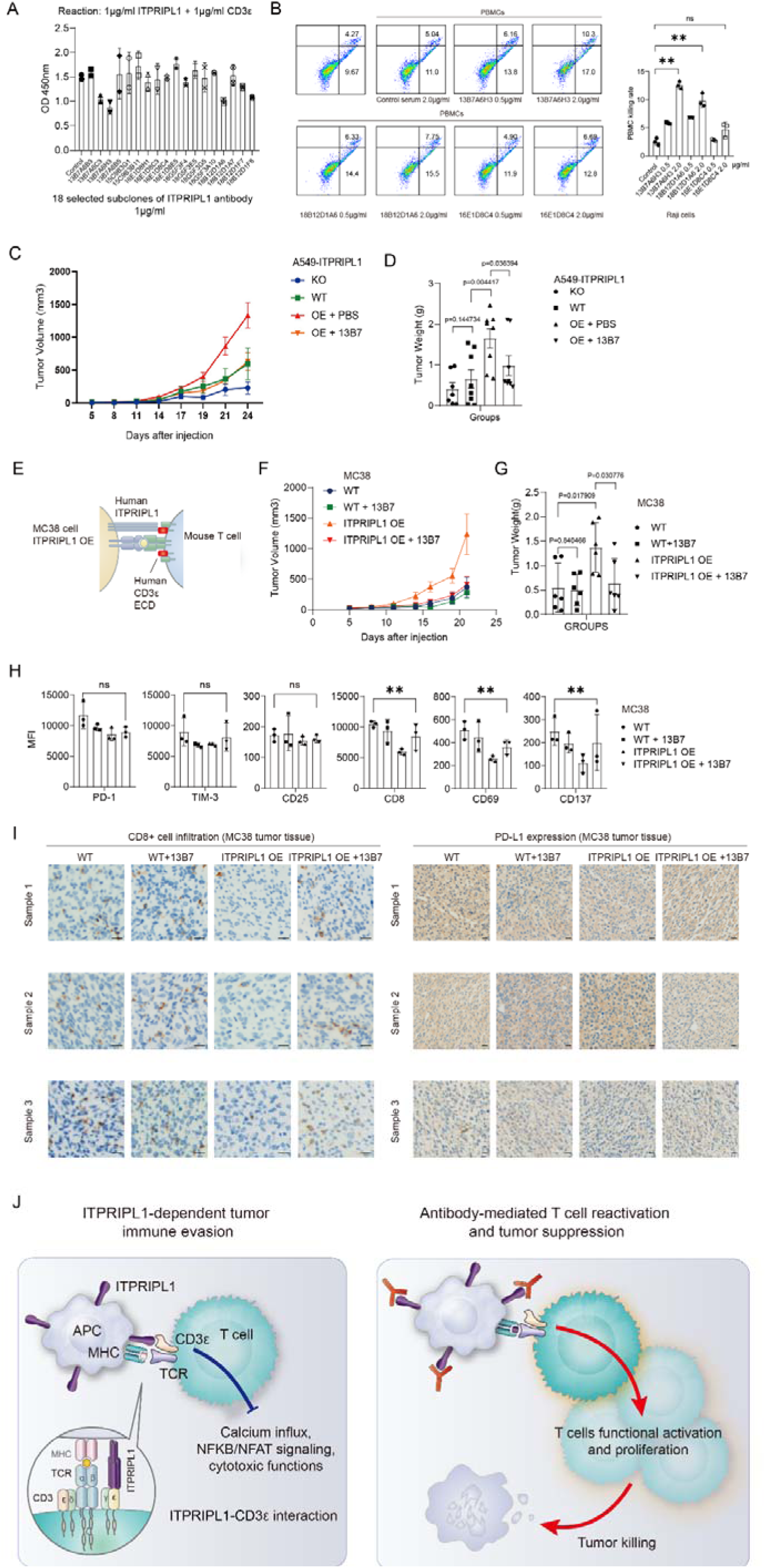
Targeting ITPRIPL1 using a monoclonal antibody de-repress T cells activation. **A**. ELISA showing the ability of 18 selected subclones to block ITPRIPL1-CD3ε interaction, with 13B7A6H3 showing the greatest effect (n=2). **B**, PBMC killing assay showing ITPRIPL1 monoclonal antibodies can promote the T cell killing functions (n=3). **C**, A549 tumor growth curves showing ITPRIPL1 promoted tumor growth while ITPRIPL1 knockout and 13B7 antibody counteracted such promotion (n=8) (FDR-adjusted P<0.05). **D**, The tumor weight of different groups (n=8) (FDR-adjusted P<0.05). **E**, Schematic of the CD3ε-e(hCD3ε)1 mice model,. **F**, MC38 tumor growth curves showing ITPRIPL1 promoted tumor growth while 13B7 antibody counteracted such promotion (n=5) (FDR-adjusted P<0.05). **G**, Bar graph of the tumor weight of the 4 experimental groups at the end of the experiment (n=6) (FDR-adjusted P<0.05). **H**, F Bar graph illustrating the mean fluorescence intensity (MFI) of a panel of immune checkpoint markers in the 4 experimental groups as in **G** by FACS. It shows obvious changes mediated by ITPRIPL1 and 13B7 antibody in CD8, CD69, CD137 in the peripheral blood (n=3). **I**, Histological evaluation of tumors from the 4 experimental groups as in **G** at 200X magnification (left panels) and 100X magnification (right panels). IHC revealed ITPRIPL1 OE decreased CD8^+^ T cell infiltration, which can be counteracted by 13B7 antibody in 200X and 100X views, but PD-L1 expression did not undergo obvious changes (n=3). **J**, Brief schematic of the possible mechanism of ITPRIPL1-CD3ε regulation promoting tumor immune evasion. Such inhibition can be counteracted by 13B7A6H3 monoclonal antibody. Data are mean ± s.d. **P<0.05, ***P<0.001, ****P<0.0001. Two-tailed Student’s t-test. Except for the tumor growth curves (± s.e.m.) and tumor weight (one-tailed Student’s t-test.).

To demonstrate the potential future use of the 13B7 monoclonal antibody in the clinic, we test its efficacy at repressing tumor growth in two orthogonal humanized mouse models. First, we use the humanized PBMC mouse model [25,26]. Briefly, this model consists of co-implanting cancer cells (A549) and PBMCs subcutaneously at 3:1 ratio after one week of adaptation to the environment followed by euthanasia at day 24 [27,28]. To evaluate the critical role of ITPRIPL1, we derived A549 cell overexpressing ITPRIPL1 (A549-ITPRIPL1 OE) and A549 cell ITPRIPL1 knockout (A549-ITPRIPL1 KO) (Supplementary Fig.3F). The group of mice injected with A549-ITPRIPL1 KO cells (+PBMCs) display reduced tumor size while tumors from the group of mice injected with A549-ITPRIPL1 OE (+PBMCs) are larger (Fig. 4C-D). The latter effect can be counteracted by specific ITPRIPL1-targeting using the monoclonal antibody 13B7 (Fig. 4C-D).

We established a mouse colorectal MC38 subcutaneous transplanted tumor models in CD3ε-e(hCD3ε)1 mouse in a second model. Importantly we tested that human ITPRIPL1 did not interact with mouse CD3ε (Supplementary Fig.4G). We also established and validated an ITPRIPL1 overexpressing MC38 cell line (MC38 ITPRIPL1 OE) (Supplementary Fig.4H). The humanized CD3ε-e(hCD3ε)1 mouse model is depicted in Fig.4E. As expected, mice injected with MC38 ITPRIPL1 OE substantially upregulate tumor growth. The 13B7 antibody treatment can rescue that growth compared to the group treated with mouse IgG or injected with the parental MC38 cells (Fig. 4F-G). To make sure the effects are due to T cell activation and not T cell exhaustion, we isolated the resulting PBMCs and evaluated the expression of a panel of T cells exhaustion markers including (i.e. the PD-1, LAG-3, TIM-3, CD25 [29-34]), and activating hallmarks CD69, CD137 [35,36] by FACS. The results revealed that ITPRIPL1 did not upregulate the expression of T cells exhaustion markers but significantly downregulated CD69 and CD137 expression, which can be partly rescued by the 13B7 antibody (Fig. 4H). Such changes in markers indicated that ITPRIPL1 directly inhibited the activation of T cells. We tested the expression of the hallmarks of effector T cells infiltration by immunohistochemistry. The results suggested that ITPRIPL1 can significantly downregulate CD8^+^ T cell infiltration, while PD-L1 expression was not substantially affected (Fig.4I). Taken together, our proof-of-principle pre-clinical studies revealed that ITPRIPL1 overexpression can directly downregulate T cell activation and promote MC38 tumor growth *in vivo*, which can be rescued by the 13B7 monoclonal antibody, paving the way to a novel line of immunotherapies.

## Discussion

The discovery of immune checkpoints had revolutionized immunotherapies. This study reports a novel immune checkpoint: the ligand ITPRIPL1 and its receptor CD3ε. Although CD3ε has not been reported to be a receptor or have any natural ligand, several antibody clones including OKT3, UCHT1, HIT3A, TRX4, and HuM291 have been found to bind CD3ε and to regulate T cell functions [30]. The OKT3 clone represents the first monoclonal antibody approved by FDA to be used as an immunosuppressant after transplantation [31]. Its mechanism of action remains unclear. Although OKT3 induces T cell proliferation *in vitro*, it depletes T cells *in vivo*, a seemingly contradictory effect shared with other clones [32-34]. The CD3ε subunit exhibits unique features in the TCR-CD3 complex in at least three aspects: 1) only CD3ε-specific antibodies can regulate T cells; 2) only CD3ε forms two different heterodimers with other subunits; and 3) the CD3ε intracellular domain plays multifaceted roles not found for other subunits [22]. Since CD3ε is believed to function as an adaptor protein transducing TCR signal into the T cell, the CD3ε antibodies are thought to function “accidentally” by creating trans-interaction with CD3ε ectodomain, which is thought to represent an unnaturally occurring scenario.

The discovery of intrinsic CD3ε ligand explains the importance and complexity of CD3ε, and further highlights CD3ε as another pole of functional regulation in addition to TCR. Hence, there are at least two different input signals to the TCR-CD3 complex: one mediated by ligation of pMHC to TCR representing an “on” signal, while the other is triggered by the ligation of ITPRIPL1 to CD3ε, delivering an “off” signal to signal one. This mechanism allows both sensitive recognition of neoantigen through TCR and secured tolerance to self-antigens with high T-cell immunogenicity, such as the antigens from spermatids in the testis.

Our data lead to reconsider the functions of CD3ε as a bona fide receptor with at least one physiological ligand, ITPRIPL1 (or may be named as CD3L1 to reflect the functions of this poorly uncharacterized gene better). Prior to this report, there was no function ascribed to ITPRIPL1. We developed a series of powerful genetic and pharmacological tools both *in vitro* and *in vivo* to demonstrate that ITPRIPL1 represses T cell activation through CD3e. We provide the proof of principle that using the 13B7 monoclonal antibody against ITPRIPL1 can serve as immunotherapy in pre-clinical mouse models. Since expression of PD-L1 is very often mutually exclusive with ITPRIPL1 and patients that are unresponsive to an anti-PD-1 therapy express high levels of ITPRIPL1, it is tempting to speculate that an anti-ITPRIPL1 therapy could be beneficial to non-responders of anti-PD-1 therapies and patients with PD-1 negative tumors.

## Materials and methods

### Tumor models

All animal experiments were performed in strict accordance with the relevant ethical guidelines, approved by the department of laboratory animal science of Fudan University and the Institutional Animal Care and Use Committee of Renji Hospital, School of Medicine, Shanghai Jiaotong University. After one week of adaptation to the environment, each individual CD3ε-e(hCD3ε)1 mouse (female, 4 weeks old) was randomized into four groups (n=6 for each group) and injected 1.5×10^6^ pre-treated mouse colorectal MC38 cells or MC38-ITPRIPL1 stable cells subcutaneously in the right flank (the establishment of MC38 stable clones as described later). Since Day5 after inoculation, the tumor sizes were recorded every 2-3 days using a vernier caliper and calculated with the formula 1/2 × A × a^2^ (A and a, respectively, denote the length and the width of the tumor). We treated the two groups of wild type MC38 colorectal mice models with 100μg mouse IgG or monoclonal 13B7A6H3 antibody respectively from the fifth day after inoculation of tumor cells every three days, four times in total. The two ITPRIPL1-overexpressing groups were treated alike. In accordance with the ethical guidelines, mice would be sacrificed once the tumor volume reached 2cm^3^ or ulcers happened. In this experiment, all the mice were sacrificed on day 23 after inoculation and the tumors were resected and weighed, and pre-treated for further experiments.

### ITPRIPL1 KO mouse models

C57BL/6J mice embryos were collected and CRISPER/Cas9 technique was applied to knockout ITPRIPL1 from the embryos. The KO status was validated by PCR. The KO embryos were then cultured and fed to breed their next generations. First-generation mosaic mice were crossed with C57BL/6J wild-type mice to obtain heterozygous mice (HE), and heterozygous mice were crossed to obtain homozygous mice (HO). We bred these mice till 8 weeks old and extracted sperm (n=3 for each group), and then sacrificed them for further experiments.

### Humanized PBMC mouse models

All animal experiments were performed in strict accordance with the relevant ethical guidelines, approved by the department of laboratory animal science of Fudan University and the Institutional Animal Care and Use Committee of Renji Hospital, School of Medicine, Shanghai Jiaotong University. After one week of adaptation to the environment, each individual NPSG mouse (female, 5 weeks old) was randomized into four groups (n=8 for each group) and injected 1×10^6^ pre-treated A549-ITPRIPL1 KO/WT/OE cells with 3.3 × 10^5^ PBMCs subcutaneously in the right flank. Since Day5 after inoculation, the tumor sizes were recorded every 2-3 days using a vernier caliper and calculated with the formula 1/2 × A × a^2^ (A and a, respectively, denote the length and the width of the tumor). We treated the one group of the A549-ITPRIPL1 OE mice models with 100μg monoclonal 13B7A6H3 antibody and the others with PBS from the fifth day after inoculation of tumor cells every three days, four times in total. In accordance with the ethical guidelines, mice would be sacrificed once the tumor volume reached 2cm^3^ or ulcers happened. In this experiment, all the mice were sacrificed on day 24 after inoculation and the tumors were resected and weighed, and pre-treated for further experiments.

### Cell Culture

All cell lines including human CRC HCT116 cells, NSCLC A549 cells, human blood cancer Jurkat, Raji cells were purchased from ATCC, mouse colorectal cancer MC38 cells and melanoma B16 cells were purchased from Kerafast, luciferase-reporter Jurkat-dual cells were purchased from Invivogen, PBMCs were purchased from Mt-bio, HEK293 cells were given from Zhigang Lu’s Lab (IBS, Fudan, Shanghai, China) and all mycoplasma free. HCT116, Jurkat, Raji and PBMCs were incubated in RPMI-1640 (Meilunbio) with 10% FBS (Gibco). A549 and MC38 cells were incubated in DMEM (Meilunbio) with 10% FBS (Gibco). Jurkat-dual cells were incubated in IMDM (Meilunbio) with 10% FBS (Gibco) and supplemental selective antibiotics (Invivogen). HEK293 cells were incubated in 293-union medium (Union-biotech). All cells were cultured at 37°C under 5% CO2.

### Antibodies and Reagents

The primary antibodies for GAPDH (KC-5G5, KANGCHEN), ITPRIPL1 (TA336137, Origene), Flag-tag (4793, CST), PD-1 (18106-1-AP, Proteintech), PD-L1 (66248-1-Ig, Proteintech), CD3-FITC (11-0032-82, Invitrogen), CD4-PE (12-0081-82, Invitrogen), CD8 (12-0041-82, Invitrogen), PD-L1-PE (12-5982-82, Invitrogen), PD-1-FITC (11-9985-82, Invitrogen), CTLA-4-PE (12-1522-82, Invitrogen), CD39-PE (12-0391-82, Invitrogen), CD73-PE (12-0731-82, Invitrogen), PD-1-APC (135209, Biolegend), CD25-APC (101909, Biolegend), TIM3-APC (134007, Biolegend), CD8-APC (140410, Biolegend), CD69-APC (104513, Biolegend), CD137-APC (106109, Biolegend) were all commercially available. Reagents involving recombinant protein CD3ε (10977-H02H; SinoBiological), CTLA-4 (CT4-H5255; Acrobiosystems), CD28 (CD8-H525a; Acrobiosystems), CD96 (TAE-H5252; Acrobiosystems), LAG-3 (LA3-H5255; Acrobiosystems), TIM-3 (TM3-H5258; Acrobiosystems), CD40 (CD0-5253; Acrobiosystems), SIRP alpha (SIA-H52A8; Acrobiosystems), GITR (GIR-H5254; Acrobiosystems), BTLA (BTA-H5255; Acrobiosystems), ICOS (ICS-H5258; Acrobiosystems), OX40 (OX0-H5255; Acrobiosystems), TIGHT (TIT-H5254; Acrobiosystems), LY86 (10242-H02H; SinoBiological), LILRB2 (14132-H02H; SinoBiological), LILRB4 (16742-H02H; SinoBiological), CD27 (CD7-H5254; Acrobiosystems), PD-1 (10377-H02H; SinoBiological), LTBR (LTR-H5251; Acrobiosystems), NRP1 (NR1-H5252; Acrobiosystems) were also purchased from the indicated suppliers.

### PCR

PCR reaction system component 1: ddH2O 14.9 µl, 10 x Taq PCR buffer 2 µl, 2.5 mM dNTP 1 µl, primer I (10pmol/µl) 0.5 µl, primer II (10pmol/µl) 0.5 µl, Taq DNA polymerase 0.1 µl, genomic DNA 1 µl. PCR reaction system component 2: ddH2O 14.9 µl, 10 x Taq PCR buffer 2 µl, 2.5 mM dNTP 1 µl, primer III (10pmol/µl) 0.5 µl, primer IV (10pmol/µl) 0.5 µl, Taq DNA polymerase 0.1 µl, genomic DNA 1 µl. (Primer I: CCCTGAATTCTGCAGGCACT; Primer II: TCCAGGTCCAGGGGAACTAA; Primer III: AATTACTGGGTGAAGGCCG; Primer IV: GTAGGAGGGTGGGGATCAGT; all 5’ to 3’). Step 1: 95°C, 5min; Step 2: 95°C, 15sec; Step 3: 60°C, 15sec; Step 4: 72°C, 2min; (repeat steps 2-4 for 35 cycles); Step 5: 72°C, 5min; Step 6: 12°C, hold.

### Mass spectrum

Two groups of PBMCs (5×10^6^ cells/ml) were stimulated with 1μg/ml anti-CD3/CD28 antibodies (Invitrogen) at 37°C under 5% CO2 for 18 hours in RPMI-1640 medium. One group was treated with ITPRIPL1-Fc protein (2μg/ml) and the other with identical volume of PBS (Applied Cells, Inc.). After 18 hours of treatment, we collected the samples and washed them with PBS twice, and lysed the cells with RIPA lysis buffer (Beyotime) containing 1% cocktail of proteinase and phosphatase inhibitor and PMSF (KANGCHEN) on ice for 30 minutes. The cell lysates were collected and centrifuged (12,000 rpm, 15 min, 4°C). We detected the protein concentration by BCA Protein Assay Kit (Meilunbio) and selected 100μg total protein for each group to undergo mass spectrum (MALDI-TOF). Each sample was analyzed three times. We performed gene set enrichment analysis (GSEA 4.1.0) and summarized the enrichment map.

### Luciferase assay

The Jurkat-dual cells were resuspended in test medium IMDM (Meilunbio) +10%FBS (Gibco) to 2×10^6^/ml. We then added 180μl of cell suspension per well of a flat-bottom 96-well plate (Costar) with indicated concentrations of ITPRIPL1 protein. For the groups that required activation, 50μg/ml ConA (Aladdin) for Jurkat-dual cells was added simultaneously. To coat protein on beads, we first placed ITPRIPL1 protein with 6x-His antibody (Invitrogen) first and slowly rotated at room temperature for 1 hour, and added protein A beads (Smart-Lifesciences) and slowly rotated at room temperature for another 1 hour. We incubated the plate at 37°C under 5% CO2 for 18 hours. We then added 20μl sample per well into a 96-well white plate (Costar), then 50μl Quanti-luc solution (Invivogen) into each well. The plate was placed in SpectraMax i3x and proceed measurement immediately. The whole process was protected from light.

### Protein-protein binding ELISA

The protein was diluted in coating buffer (Solarbio) to indicated concentrations and added 100 μl per well into a 96-well ELISA plate (Costar). We placed the plate at 4°C overnight. We washed the plate with PBST (Applied Cells, Inc.) five times and added 100 μl 5% BSA (VWR Life Sciences) into each well and incubated at 37°C for 90 minutes. We repeated washing and added 100 μl protein into each well and incubated at 37°C for 60 minutes. We repeated washing and added 100 μl associated secondary antibodies conjugated with HRP into each well and incubated at 37°C for 30 minutes. After 5 times PBST washing, we added 100 μl TMB buffer (Invitrogen) into each well and incubated at 37°C for 15 minutes, and added 50 μl stop buffer (Abcam) to terminate the reaction. The plate was placed in SpectraMax i3x and read at 450 nm.

### FACS

The cells (2×10^6^/ml) were added 100 μl per well a flat-bottom 96-well plate (Costar) with indicated concentrations of protein. We incubated the plate at 37°C under 5% CO2 for 30 minutes. After incubation, we washed the samples with flow cytometry staining buffer (Invitrogen) three times. Then we diluted the fluorescent antibodies at suggested concentrations according to the suppliers in staining buffer and added 200 μl to each sample. We incubated them at room temperature for 30 minutes, protected from light. Next, after being washed with staining buffer three times, we transferred the samples into single tubes (Falcon) and analyzed them by MACSQuant16 (Miltenyi). We applied FlowJo V10 to analyze the data.

### ITPRIPL1 protein expression and purification

The gene ITPRIPL1 was synthesized (General Biosystems (Anhui) Co., Ltd) and cloned into modified pcDNA3.1 vector. The plasmid was transfected into HEK293 cells using PEI. After culture at 37°C under 5% CO2 for 6 days, cells were collected and lysed by 1XPBS (pH 7.2-7.4), 0.5% CHAPS at 4°C for 30 min, and the insoluble fraction was removed by centrifugation at 30,000 x g for 30 min. Supernatants were incubated with protein A-agarose for 1-2 h and washed extensively. The ITPRIPL1 protein was eluted by 0.1M glycine (pH 3.0) and neutralized with 1M Tris-HCl (pH8.5), and then concentrated to 1mg/ml and stored at -80°C.

### Plasmids construction

The pcDNA3.1-Flag-ITPRIPL1 vector was established by inserting synthesized cDNA encoding Flag tag and ITPRIPL1 (shanghai Generay Biotech Co., Ltd) into pcDNA3.1 vector using EcoRI/XhoI MCS. The pcDNA3.1-HA-CD3ε was purchased from Generay, constructed by inserting synthesized cDNA encoding HA tag and CD3ε (human) into the pcDNA3.1 vector, using the EcoRI/XhoI multiple cloning sites (MCS). The pcDNA3.1-Flag/Fc-ITPRIPL1 RBD/RBD2/RBD3 plasmids were purchased from General Biol, generated by inserting synthesized cDNA encoding Flag tag or a tag containing the hinge region, CH2 region and CH3 region of Fc with ITPRIPL1 RBD/RBD2/RBD3 sequence into the pcDNA3.1 vector, using the EcoRI/XhoI MCS. All vectors were checked by sequencing and western bolt with specific antibodies in which the observed molecular weights were in concordance with the predicted molecular weights.

### Transfection of plasmids

Tumor cells were seeded in 6-well plates to reach a density of 50∼70% at the time of transfection. 24 hours later, transfection was performed using 1.5μg plasmid together with 4.5μl FuGENE HD (Promega) and 100μl Opti-MEM per well according to the manufacturer’s guidance. The negative control in each experiment was cells mock-transfected with empty control vector.

### Establishment of ITPRIPL1/CD3ε stable cells

We chose human CRC HCT116 cells to establish ITPRIPL1 stable cell strains. Firstly, we transfected ectopic Flag-tagged ITPRIPL1 or HA-tagged CD3ε into the HCT116 cells following the procedures described above. We also transfected a blank vector control and an empty control, respectively. After approximately two-week incubation in RPMI-1640 (Gibco) supplemented with 1000 μg/ml G418 (Gibco BRL) with refreshing the medium every 2-3 days, the single colonies were picked and verified by immunoblots. Given the results of the overexpression of ITPRIPL1/CD3ε determined by western blot, we chose a colony with highest ITPRIPL1/CD3ε expression and generated the ITPRIPL1/CD3ε stable cells.

### Lentivirus transduction

The lentiviruses for CD3ε silencing and CD3ε mutants were purchased from Genepharma, and the Cas9 and ITPRIPL1 sgRNA lentiviruses were purchased from Genomeditech. The sequences of lentiviral sgRNA and shRNA are summarized in Supplementary Table 2. HCT116 cells were seeded in a 6-well plate, and were infected with Cas9 lentivirus and blank vector in RPMI-1640 complete medium supplemented with 5 μg/ml polybrene to construct HCT116-Cas9 cell line. After 24h incubation, the medium was refreshed with RPMI-1640 complete medium containing 10 μg/ml blasticidin (Invivogen). The medium was refreshed every 2-3 days for two weeks and identified the transfection efficiency by immunoblot. Then we transfect the HCT116-Cas9 cell line with lentivirus containing different sgRNAs targeting ITPRIPL1 and blank vector, added 1 μg/ml puromycin (Invivogen) to select out the cell line with best ITPRIPL1 knock out effect, confirmed by immunoblot. Jurkat cells were seeded in a 6-well plate and infected with CD3ε-specific shRNA lentivirus, supplemented with 5 μg/ml polybrene. After similar refreshment of medium and 1 μg/ml puromycin for selection, we tested the knockdown effect by immunoblot and selected the best knockdown group. The CD3ε knockdown Jurkat cells were infected with CD3ε-ΔPRS-Flag or CD3ε-K76T-Flag lentivirus. The following procedures were similar, with 600 μg/ml G418 (Gibco BRL) for selection. The expression of the CD3ε mutants were tested by immunoblot. The expression of ITPRIPL1 and CD3ε was also examined by RT-PCR and immunoblots before use.

### Immunoblotting

Cells were lysed with RIPA buffer (Beyotime) supplemented with 1% proteinase and phosphatase inhibitors cocktail (ThermoFisher Scientific). The collected cell lysates were centrifuged for 15min at 12000rpm (4°C). The supernatant was reserved and the protein concentration was determined by BCA Protein Assay Kit (ThermoFisher Scientific). 5×SDS-PAGE loading buffer (Applied Cells, Inc.) was diluted to 1× with protein sample and heating at 100°C for 8 min. The protein extracts were subjected to appropriate concentrations of SDS–PAGE for electrophoresis and transferred to PVDF membranes (Bio-Rad). Membranes were blocked with 5% bovine serum albumin (ThermoFisher Scientific) for one hour at room temperature, and then incubated with the primary antibodies overnight at four degrees. Membranes were incubated with secondary HRP-conjugated antibodies (KANGCHEN) at room temperature for one hour. Before and after the incubation, the membranes were washed five times with TBST and then examined by ChemiDoc imaging system (Bio-Rad).

### Immunohistochemistry

The MC38 tumor specimens were incubated with antibodies against PD-L1 (Cat17952-1-AP, 1:200; Proteintech) and CD8α (Cat98941, 1:200; Cell Signaling Technology). After three times PBS washing, the tissues were then incubated with a biotin-conjugated secondary antibody, followed by avidin-biotin–peroxidase complex. Visualization was performed using aminoethyl carbazole chromogen.

### Mice tissue HE stain

The ITPRIPL1 WT/HE/HO mice were obtained and sacrificed as described above. We extracted the heart, liver, spleen, brain, salivary gland, pancreas from them and fixed with 4% paraform aldehyde. We washed the fixed tissue and dehydrated with increasing concentrations of ethanol. Then we waxdip and embed the tissue to make paraffin slides. The slides were dewaxed and hydrated by dimethylbenzene and decreasing concentrations of ethanol. The antigen retrieval was done by citrate. We then followed the instruction guide provided by Solarbio HE stain agents. The tissues were observed by microscope.

### Mice sperm motility observation

The ITPRIPL1 WT/HE/HO mice were obtained as described above. After placing specific electronic device into the rectum of each mouse, we gradually increased voltage until ejaculation. The sperm was collected, spread evenly and observed directly under microscope. The sperm motility was evaluated according to WHO laboratory manual for the examination and processing of human semen (fifth edition, 2010).

### T-cell killing assay

The PBMCs were stimulated with 1μg/mL CD3 antibody (317303, BioLegend), 1μg/mL CD28 antibody (302913, BioLegend), and 10 ng/mL IL-2 (589102, BioLegend) in RPMI-1640 complete medium for 24 hours before killing to differentiate into T cells. T cells and the killing target cells were counted and added 2×10^5^ to each well of a 24-well plate in 1:1 ratio. After 6 hours of killing, cells were collected and washed with flow cytometry staining buffer (Invitrogen) three times. We stained the mixture with CD45-APC antibody (17-0459-42, Invitrogen) for 30 minutes at room temperature. Then we washed the samples with flow cytometry staining buffer three times, and stained them with Annexin-V-FITC and PI (Beyotime) for 15 minutes at room temperature. We transferred the samples into single tubes (Falcon) and analyzed them by MACSQuant16 (Miltenyi). We applied FlowJo V10 to analyze the data.

### OCTET binding kinetics assay

The ITPRIPL1 protein was His-tagged and CD3ε protein was Fc-tagged. We applied His-tag probe and set ITPRIPL1 protein as the antigen while CD3ε protein as the antibody. The whole antigen-antibody affinity binding kinetics assay was performed according to the instruction guide of Gator (SNGC00070, ProbeLife, Lnc.), and data were collected and analyzed by Gator software.

### Identification of intracellular calcium flux

After treatment for 24h, cells were stained with Fluo-8 AM (ab142773) (100 μl, 4 μM) in HHBS at 37 °C under 5% CO2 for 1h, protected from light. Then after being washed twice with HHBS, cells were observed with a fluorescence microscope.

### Data availability

The authors declare that all data supporting the findings of this study are available within the paper and its Supplementary Information files.

## Acknowledgements

This work was supported by National Natural Science Foundation of China (No: 82030104, 81874050, 81572326), Basic Research Projects of Shanghai Science and Technology Innovation Action Plan (20JC1410700); National Key R & D Program of China (2016YFC0906002, 2016YFC0906002), Tang Scholar (XJ), and Startup Research Funding of Fudan University.

## Author contributions

SD, YTW, YGW performed experiments and analyzed data. SD, JBP and XJ wrote the paper. XJ conceived the study and provided resources for the experimental researches.

