## Supplementary Information for "The ITPRIPL1- CD3ε axis: a novel immune checkpoint controlling T cells activation"

**Supplementary Tables**

**Table.S1. T cell all significant gene sets.**

| EnrichmentMap::Formatted_name | EnrichmentMap::Genes | EnrichmentMap::gs_size | EnrichmentMap::pvalue (gene_sets) | name |
| --- | --- | --- | --- | --- |
| GSE21360_NAIVE_VS_ QUATERNARY_ MEMORY_CD8_TCELL_ DN | FAM98A\|ARMCX3\|CDC73\|MIS12\|NGLY1\|ELF1\|DDX47\|SP100\|IFITM2\|CD38\|TRIM21\|RBM22\|ANKFY1\|EIF2AK2\|MOB1A\|SAMD9\|CRK\|SERPINB9\|MX1\|OAS3\|ADPGK\|CUL4A\|PPP2R2A\|LAMP3\|DCP1A\|EIF5\|EIF4A1\|TRAPPC4\|BST2\|BNIP1\|ZNF148\|PSMD11\|POU2F1\|ARPC1A\|MYCBP2\|RIPK1\|TREX1\|NIP7\|RSRC2\|PRRC1\|POGLUT1\|DDX18\|LAP3\|CASP1\|N4BP1\|NOC3L\|NIPBL\|NADK\|IFIH1\|ISG20\|CHMP5\|LNPEP\|NMI\|ISG15\|TRAFD1\|OASL\|NFKBIE\|TNPO2\|EIF2S1\|POLG\|DDX60\|IFIT1\|ADAR\|PLSCR1\|TRIM22\|TLR3\|AARSD1\|IFIT2\|UBE2L6\|TAP1\|DDX58\|MX2\|IRF9\|MRTO4\|LGALS8\|IFIT5\|PSMA6\|GOSR1\|NBN\|STAT2\|GTF2B\|LARP1\|IFI35\|DENND1A\|OAS2\|PML\|BZW2\|EHD4\|CASP4 | 89 | 0.01 | GSE21360_NAIVE_VS_QUATERNARY_MEMORY_CD8_TCELL_DN |
| GSE19888_CTRL_VS_ TCELL_MEMBRANES_ ACT_MAST_CELL_ PRETREAT_A3R_INH_ DN | APOD\|NFKB1\|GZMA\|TOR3A\|CD40\|MPEG1\|TNFAIP2\|FPR2\|DTX3L\|EIF4E3\|PTPN6\|LACC1\|IKZF1\|AIDA\|TRIM21\|SIRPB1\|ANKFY1\|EIF2AK2\|DCK\|SAMHD1\|TAPBP\|SERPINB9\|IL4I1\|SLFN5\|CMPK2\|CCR7\|SAMD9L\|PARP14\|SCLY\|HK1\|SH2D1A\|BST2\|GZMB\|ARHGAP30\|NPC2\|PIK3R5\|TOR1AIP1\|TUBB6\|CARM1\|CNP\|HLA-E\|N4BP1\|GMPPB\|PARP9\|TOR1AIP2\|APOL2\|IFIH1\|USP11\|NMI\|GCH1\|GLA\|TRIM25\|PIK3AP1\|DDX60\|IFIT1\|ADAR\|ARF4\|TLR3\|SEMA4D\|IFIT2\|INPP1\|TLR2\|BCL3\|IRF9\|ADA\|SLAMF7\|CD274\|UBA7\|SF3A2\|RHOH\|VDAC2\|CLIC4\|GNB4\|NAAA\|STAT2\|CASP9\|LARP1\|IDO1\|RAC1\|C8ORF33\|STAT5B\|VCAM1\|EVI2B\|PTK2B\|GRN\|ITGB7\|OAS2\|ATG16L1\|ITGAL\|RFX1\|CD300LF\|ETV6 | 92 | 0.01 | GSE19888_CTRL_VS_TCELL_MEMBRANES_ACT_MAST_CELL_PRETREAT_A3R_INH_DN |
| GSE21360_NAIVE_VS_ QUATERNARY_ MEMORY_CD8_TCELL_ UP | CDC73\|CMAS\|MIS12\|ELF1\|SP100\|IFITM2\|CD38\|UTP14C\|ARFIP1\|TRIM21\|RBM22\|EIF2AK2\|SAMD9\|CRK\|MX1\|OAS3\|PCID2\|CD48\|XRCC4\|GLS\|WDR44\|PPP2R2A\|HERC4\|LAMP3\|DCP1A\|EIF5\|IMPAD1\|EIF4A1\|TRAPPC4\|TMED5\|BST2\|SFSWAP\|AGA\|RAP1B\|ARMC8\|RIPK1\|IDH3A\|ARFGEF2\|PRRC1\|ACP1\|LAP3\|N4BP1\|NOC3L\|PIGK\|TMEM9B\|TCP1\|NADK\|USP16\|VCPIP1\|IFIH1\|ISG20\|CHMP5\|LNPEP\|ORC4\|NMI\|ISG15\|OASL\|TRIM25\|DDX60\|IFIT1\|PLSCR1\|ADAR\|TRIM22\|AARSD1\|IFIT2\|DCTN4\|MX2\|DDX58\|SNX2\|IRF9\|RBM7\|TOP1\|IFIT5\|STAT2\|TMEM33\|QKI\|GTF2B\|LARP1\|IFI35\|SERPINB8\|STAMBP\|SYNJ1\|OAS2\|EXOSC2\|SAR1B\|BZW2\|EHD4\|IPO8 | 88 | 0.01 | GSE21360_NAIVE_VS_QUATERNARY_MEMORY_CD8_TCELL_UP |
| GSE37533_PPARG1_ FOXP3_VS_FOXP3_ TRANSDUCED_CD4_ TCELL_ PIOGLITAZONE_ TREATED_UP | B2M\|HK2\|ELF1\|SCAMP1\|IFITM2\|EIF2AK2\|SAMHD1\|STK24\|SAMD9\|RAB9A\|MX1\|OAS3\|NT5C2\|COMMD8\|PSEN1\|PPP2R2A\|FMR1\|EFR3A\|NAPA\|BCL2L13\|BST2\|CBR3\|UNC93B1\|MYCBP2\|NR3C1\|RABAC1\|HLA-C\|TREX1\|PDXK\|SLC22A18\|OGFR\|CASP1\|CNP\|N4BP1\|IFIH1\|LGMN\|ISG20\|LNPEP\|APOBEC3G\|NMI\|CTNNBL1\|ISG15\|TARBP1\|OASL\|TRADD\|GCH1\|SPTLC2\|TRIM25\|MYD88\|PGAM1\|KAT2B\|DDX60\|IFIT1\|PLSCR1\|ADAR\|PARP4\|IFIT2\|UBE2L6\|MEF2C\|MX2\|DDX58\|USP25\|GYG1\|PSMB9\|UBA7\|P2RX1\|IFI16\|SLC15A3\|SMARCA5\|IFIT5\|LGALS8\|LGALS9\|NBN\|RBCK1\|PSMB8\|PRKD2\|TRIM26\|IFI35\|TOR1B\|BAG1\|NFYB\|CD47\|OAS2\|PML\|TRANK1\|FHL3\|HLA-F\|EHD4\|SUGP1\|EML2 | 90 | 0.01 | GSE37533_PPARG1_FOXP3_VS_FOXP3_TRANSDUCED_CD4_TCELL_PIOGLITAZONE_TREATED_UP |
| GSE37533_PPARG1_ FOXP3_VS_PPARG2_ FOXP3_TRANSDUCED_ CD4_TCELL_ PIOGLITAZONE_ TREATED_DN | SNTB2\|SNRK\|TNFAIP2\|GBP2\|FKBP5\|CASP10\|OAS3\|VAMP5\|FES\|BCL2L13\|SDAD1\|ACSL5\|HLA-E\|ZC3HAV1\|IRF2\|IFIH1\|CYLD\|NMI\|TRAFD1\|GCLM\|ISG15\|HLA-A\|ENY2\|GCH1\|BACH1\|USP15\|MYD88\|DDX60\|DNAJB1\|ADAR\|IFIT1\|TRIM22\|IFIT2\|UBE2L6\|TAP1\|DDX58\|IRF9\|BCL3\|UBA7\|PSMB9\|CLIC2\|ATP2A2\|IFIT5\|LGALS9\|BAK1\|GBP1\|STAT2\|PSMB8\|IFI35\|IDO1\|AKAP8\|TAP2\|GOLM1\|CD47\|SETX\|PML\|OAS2\|TRANK1\|PSME2\|SP100\|TRIM21\|STAT1\|EIF2AK2\|SAMHD1\|MX1\|DDX23\|CBR3\|DYNLT1\|STK3\|RIPK1\|ATP6V1B2\|LAP3\|CALCOCO2\|BTN3A3\|OGFR\|IFI30\|CYTH1\|PSME1\|CASP1\|PSMB10\|DNAJA1\|PPA1\|PSMA4\|APOL2\|APOL3\|MED21\|DNAJA2\|PLSCR1\|TLR3\|NDUFA9\|MALT1\|SLC15A3\|RIPK2\|CASP7\|RAB27A\|TANK\|AHCYL2\|CTDSP2\|BAZ1A\|UBXN4\|HLA-F | 101 | 0.01 | GSE37533_PPARG1_FOXP3_VS_PPARG2_FOXP3_TRANSDUCED_CD4_TCELL_PIOGLITAZONE_TREATED_DN |
| GSE26890_CXCR1_ NEG_VS_POS_ EFFECTOR_CD8_ TCELL_UP | CPSF2\|TOR3A\|CSDE1\|MPEG1\|ELF1\|OGFRL1\|GTF3C2\|DTX3L\|CD86\|MLEC\|NCEH1\|GBP2\|ZEB2\|G3BP2\|TRIM21\|AIDA\|STAT1\|ZC3H11A\|EIF2AK2\|SAMHD1\|PNPT1\|IDI1\|SERPINB9\|RNASEL\|OAS3\|SLFN5\|CMPK2\|SAMD9L\|PARP14\|LPXN\|CCDC25\|RNPS1\|DEK\|SPATA13\|MTDH\|LSS\|PABPC1\|MRPL22\|CAMKK2\|CYBB\|KHSRP\|KBTBD2\|NIT2\|ANKLE2\|PTGES3\|HLA-E\|DNAJC13\|PPA1\|EXOC4\|VCPIP1\|DAXX\|PARP9\|PURA\|IFIH1\|ISG20\|TRAFD1\|ISG15\|TM9SF1\|OASL\|CCNL1\|RNF213\|CNOT3\|DDX60\|NAA20\|IFIT1\|EHD3\|TLR3\|MGST2\|IFIT2\|INPP1\|MX2\|DDX58\|LGALS3BP\|IRF9\|USP25\|MRPL1\|CD274\|SCO1\|NAMPT\|DPP4\|ITCH\|LARP1\|CAPRIN1\|LYN\|PI4KB\|OXSR1\|CD47\|OAS2\|PML\|BAZ1A | 90 | 0.01 | GSE26890_CXCR1_NEG_VS_POS_EFFECTOR_CD8_TCELL_UP |
| GSE2770_TGFB_AND_ IL4_ACT_VS_ACT_CD4_ TCELL_2H_DN | ELF1\|GTPBP1\|DTX3L\|SP100\|SH3GLB1\|IFITM2\|STOM\|CDS2\|CD38\|TRIM21\|EIF2AK2\|SAMD9\|OAS3\|WDFY1\|SAMD9L\|AP1G2\|FBXO7\|ARHGAP27\|NAPA\|LAMP3\|BST2\|UNC93B1\|SMCHD1\|CAPN2\|DYNLT1\|UBE2F\|TREX1\|SNW1\|LAP3\|CALCOCO2\|DRAP1\|TRIM56\|PARP9\|BLVRA\|IRF2\|IFIH1\|GIMAP4\|GSTK1\|CHMP5\|USP11\|ISG20\|NMI\|RRAGC\|TRAFD1\|ISG15\|CLIC1\|OASL\|RHEB\|SPTLC2\|GCH1\|TRIM25\|MYD88\|RAB8A\|FBXO6\|DDX60\|PARP10\|PLSCR1\|ARL8A\|MNDA\|NRBP1\|NCSTN\|PARP4\|DCTN4\|IFIT2\|UBE2L6\|C19ORF25\|MX2\|DDX58\|SAP30BP\|IRF9\|USP25\|IFI16\|RBM7\|ZNFX1\|ELMO2\|NUB1\|LGALS9\|IFIT5\|RP2\|STAT2\|OPTN\|TRIM26\|GTF2B\|TRIP4\|IFI35\|TOR1B\|CNDP2\|GNG5\|OAS2\|CCNK\|MLKL | 91 | 0.01 | GSE2770_TGFB_AND_IL4_ACT_VS_ACT_CD4_TCELL_2H_DN |
| GSE37533_PPARG2_ FOXP3_VS_FOXP3_ TRANSDUCED_CD4_ TCELL_DN | CASP10\|SAMD9\|OAS3\|ACOT9\|FES\|NAPA\|BCL2L13\|ACSL5\|HLA-C\|TREX1\|CNP\|N4BP1\|IRF2\|IFIH1\|ISG20\|CYLD\|CTNNBL1\|NMI\|ISG15\|TRAFD1\|HLA-A\|OASL\|SPTLC2\|GCH1\|MYD88\|STARD5\|DDX60\|ADAR\|TYMP\|IFIT1\|TRIM22\|RHBDF2\|PCF11\|IFIT2\|UBE2L6\|TAP1\|MX2\|DDX58\|USP25\|IRF9\|UBA7\|PSMB9\|RBM7\|MCL1\|LGALS8\|IFIT5\|LGALS9\|BAK1\|GBP1\|STAT2\|PSMB8\|PRKD2\|ENO3\|TRIM26\|GTF2B\|IDO1\|IFI35\|BAG1\|TAP2\|GOLM1\|SETX\|CD47\|PML\|OAS2\|TRANK1\|FHL3\|PSME2\|FAM20B\|ELF1\|SP100\|TRIM21\|STAT1\|EIF2AK2\|SAMHD1\|GSDMD\|RAB9A\|GMDS\|MX1\|CBR3\|RIPK1\|LAP3\|CALCOCO2\|OGFR\|BTN3A3\|IFI30\|PSME1\|CASP1\|PSMB10\|DNAJA1\|APOL2\|APOBEC3G\|APOL3\|PLSCR1\|TLR3\|NDUFA9\|MEF2C\|MALT1\|SLC15A3\|RIPK2\|CASP7\|PSMA6\|RPS18\|HLA-F | 103 | 0.01 | GSE37533_PPARG2_FOXP3_VS_FOXP3_TRANSDUCED_CD4_TCELL_DN |
| GSE37534_ UNTREATED_VS_ PIOGLITAZONE_ TREATED_CD4_TCELL_ PPARG1_AND_FOXP3_ TRASDUCED_DN | HK2\|CARHSP1\|STT3A\|ELF1\|CADM1\|SIGIRR\|IFITM2\|TRIM21\|STAT1\|EIF2AK2\|GSDMD\|MX1\|OAS3\|SYNRG\|TXNIP\|NAGK\|BRCC3\|VAMP5\|COASY\|LARS2\|NAPA\|FMOD\|DCP1A\|BCL2L13\|SLC25A40\|BST2\|TRPV2\|TMBIM1\|CYB5R3\|P2RX4\|ACAT2\|HLA-C\|TREX1\|NDRG1\|OGFR\|IFI30\|ABCF3\|CASP1\|PSMB10\|HLA-E\|COX4I1\|APOL2\|BCAT1\|ISG20\|CTNNBL1\|NMI\|TRAFD1\|ISG15\|OASL\|SNRPD2\|GCH1\|SPTLC2\|TRIM25\|MYD88\|STARD5\|IFIT1\|PLSCR1\|CBR1\|TLR3\|IFIT2\|UBE2L6\|TAP1\|COX17\|DDX58\|MX2\|CPT1A\|ERAP1\|IRF9\|PSMB9\|EZR\|SLC15A3\|RIPK2\|LGALS8\|GBP1\|RBCK1\|EXOSC9\|NBN\|STAT2\|PSMB8\|DPP4\|OPTN\|MRPL17\|PRKD2\|CALR\|TRIM26\|RPL13\|IFI35\|BAG1\|TAP2\|OAS2\|PML\|PTPRA\|SRGAP2\|HLA-F\|EHD4\|PSME2 | 96 | 0.01 | GSE37534_UNTREATED_VS_PIOGLITAZONE_TREATED_CD4_TCELL_PPARG1_AND_FOXP3_TRASDUCED_DN |
| GSE33424_CD161_ INT_VS_NEG_CD8_ TCELL_UP | TOR3A\|CSDE1\|SLC25A12\|MPEG1\|ELF1\|DTX3L\|CD86\|GBP2\|ZEB2\|SCAMP2\|TRIM21\|STAT1\|EIF2AK2\|SAMHD1\|TAPBP\|PNPT1\|RNASEL\|MX1\|OAS3\|SLFN5\|CMPK2\|PRMT5\|C7ORF50\|SAMD9L\|PARP14\|LPXN\|C19ORF12\|PFKFB3\|LY86\|PABPC1\|PSMD11\|CYBB\|LAP3\|NIT2\|NUTF2\|SMARCAL1\|DNAJC13\|HLA-E\|N4BP1\|PPA1\|CCT5\|PARP9\|DAXX\|TOR1AIP2\|IFIH1\|ISG20\|NMI\|ISG15\|TRAFD1\|OASL\|CCNL1\|RNF213\|CHRAC1\|PGD\|NMRAL1\|RPS2\|DDX60\|IFIT1\|GART\|EHD3\|TLR3\|DNAJC2\|MGST2\|IFIT2\|INPP1\|MX2\|DDX58\|IRF9\|HCK\|LGALS3BP\|CD274\|MRPL1\|NAMPT\|ASNS\|MTHFD1\|ST13\|STAT2\|HSPA5\|DPP4\|LARP1\|NOP58\|IFI35\|CD47\|OAS2\|PML\|BAZ1A\|GNL3\|CASP4\|CD300LF\|ETV6 | 90 | 0.01 | GSE33424_CD161_INT_VS_NEG_CD8_TCELL_UP |
| GSE19888_ ADENOSINE_A3R_INH_ PRETREAT_AND_ACT_ BY_A3R_VS_TCELL_ MEMBRANES_ACT_ MAST_CELL_UP | CD40\|TNFAIP2\|CD180\|TIMP1\|GBP2\|SIRPB1\|SERPINB9\|OAS3\|CMPK2\|MAP1S\|SAMD9L\|PARP14\|TFEC\|PTPRC\|GZMB\|HK3\|STX7\|TUBB6\|HLA-E\|PARP9\|SELL\|IFIH1\|LAIR1\|ISG20\|GIMAP4\|NMI\|ISG15\|TRAFD1\|OASL\|GCH1\|GLA\|PIK3AP1\|TRIM25\|ADAR\|IFIT1\|IFIT2\|TAP1\|UBE2L6\|MX2\|DDX58\|IRF9\|PSMB9\|UBA7\|SLAMF7\|CCL5\|GNB4\|ZNFX1\|NAMPT\|STAT2\|PSMB8\|P4HA1\|BID\|IFI35\|LYN\|PML\|OAS2\|PNP\|CASP4\|BST1\|PSME2\|MLKL\|TOR3A\|GIMAP7\|MPEG1\|DTX3L\|EIF4E3\|AOAH\|LACC1\|AIF1\|SAMSN1\|IKZF1\|LCP2\|TRIM21\|AIDA\|STAT1\|EIF2AK2\|SAMHD1\|DCK\|TAPBP\|GSDMD\|RNF31\|MX1\|DPYSL2\|SLFN5\|BST2\|VWA5A\|RAP1B\|TLR6\|NPC2\|PSME1\|CASP1\|PSMB10\|DAXX\|TOR1AIP2\|BTK\|RNF213\|TAPBPL\|TLR3\|INPP1\|HCK\|CD274\|TCIRG1\|NAAA\|CASP8\|RNASE6\|CCR5\|FCGR3A | 107 | 0.01 | GSE19888_ADENOSINE_A3R_INH_PRETREAT_AND_ACT_BY_A3R_VS_TCELL_MEMBRANES_ACT_MAST_CELL_UP |
| GSE37533_PPARG1_ FOXP3_VS_FOXP3_ TRANSDUCED_CD4_ TCELL_DN | GBP2\|CASP10\|SAMD9\|OAS3\|NAPA\|BCL2L13\|TMBIM1\|HLA-C\|TREX1\|CNP\|N4BP1\|HLA-E\|ZC3HAV1\|IRF2\|IFIH1\|LGMN\|ISG20\|CYLD\|ORC4\|NMI\|CTNNBL1\|TRAFD1\|ISG15\|HLA-A\|OASL\|GCH1\|SPTLC2\|MYD88\|USP15\|DDX60\|IFIT1\|ADAR\|TRIM22\|RHBDF2\|PARP4\|IFIT2\|UBE2L6\|TAP1\|DDX58\|MX2\|IRF9\|USP25\|UBA7\|PSMB9\|RBM7\|IFIT5\|LGALS9\|LGALS8\|BAK1\|GBP1\|RBCK1\|STAT2\|PSMB8\|PRKD2\|TRIM26\|GTF2B\|IDO1\|IFI35\|BAG1\|TAP2\|GOLM1\|CD47\|PML\|OAS2\|TRANK1\|FHL3\|EHD4\|PSME2\|ELF1\|GTPBP1\|SP100\|IFITM2\|TRIM21\|STAT1\|ANKFY1\|EIF2AK2\|SAMHD1\|RNF31\|GSDMD\|RAB9A\|MX1\|CBR3\|BST2\|DYNLT1\|MYCBP2\|RIPK1\|RABAC1\|LAP3\|CALCOCO2\|OGFR\|BTN3A3\|IFI30\|PSME1\|CASP1\|PSMB10\|PPA1\|DNAJA1\|APOL2\|MAX\|APOBEC3G\|APOL3\|PLSCR1\|TLR3\|NDUFA9\|MEF2C\|MALT1\|SLC15A3\|RIPK2\|CASP7\|IFI16\|NBN\|OPTN\|DNPEP\|NFYB\|BAZ1A\|HLA-F | 116 | 0.01 | GSE37533_PPARG1_FOXP3_VS_FOXP3_TRANSDUCED_CD4_TCELL_DN |
| GSE19888_ ADENOSINE_A3R_INH_ VS_TCELL_ MEMBRANES_ACT_ MAST_CELL_UP | ATM\|CD40\|TNFAIP2\|TERF2\|DTX3L\|GBP2\|TRIM21\|STAT1\|DCK\|TAPBP\|SAMHD1\|SERPINB9\|MX1\|SLFN5\|CMPK2\|RBM5\|SAMD9L\|PARP14\|RRP1B\|SH2D1A\|GZMB\|STX3\|PSME1\|CYTH1\|MVP\|HLA-E\|PSMB10\|TRIM56\|ZC3HAV1\|SGK3\|PARP9\|DAXX\|TOR1AIP2\|IFIH1\|NMI\|ISG15\|RNF213\|TRIM25\|NDUFV3\|ADAR\|IFIT1\|TLR3\|IFIT2\|SPPL2A\|TAP1\|INPP1\|UBE2L6\|MX2\|DDX58\|MTHFD2\|IRF9\|CD274\|PSMB9\|UBA7\|CLIC4\|PSMB8\|STAT2\|TRAF1\|NFATC2IP\|IFI35\|FPR1\|OAS2\|PML\|FCGR3A\|CASP4\|MYO9B\|PSME2 | 67 | 0.01 | GSE19888_ADENOSINE_A3R_INH_VS_TCELL_MEMBRANES_ACT_MAST_CELL_UP |
| GSE41978_ID2_KO_VS_ BIM_KO_KLRG1_LOW_ EFFECTOR_CD8_ TCELL_UP | CLPX\|B2M\|F2\|CSTF3\|HNRNPH2\|SNRK\|ICAM1\|SP100\|GBP2\|CXCL5\|TRIM21\|STAT1\|TAPBP\|SAMD9\|OAS3\|MX1\|PTPRC\|HK1\|DDX23\|BST2\|CYB5A\|RCN1\|STK3\|RPA2\|SOD2\|HLA-C\|MAFF\|LAP3\|CALCOCO2\|OGFR\|BTN3A3\|IFI30\|IGFLR1\|PSMB10\|HLA-E\|CTSS\|ZC3HAV1\|MET\|APOL2\|IFIH1\|MAX\|ISG20\|GSTK1\|LSM6\|NMI\|APOL3\|ISG15\|TRAFD1\|HLA-A\|GCLM\|SERPING1\|OASL\|BACH1\|MYD88\|TRIO\|C3\|DDX60\|PLSCR1\|IFIT1\|STAT3\|ADAR\|CFH\|TRIM22\|CEBPB\|TLR3\|UBE2L6\|TAP1\|DDX58\|IRF9\|LGALS3BP\|PSMB9\|RABGAP1L\|CASP7\|IFI16\|TOP1\|RAB27A\|IFIT5\|NBN\|GBP1\|RBCK1\|PSMB8\|CASP8\|GTF2B\|SYNE2\|IDO1\|IFI35\|TAP2\|CLEC2B\|KDSR\|RBM4\|CD47\|PML\|TRAT1\|PSME2 | 94 | 0.01 | GSE41978_ID2_KO_VS_BIM_KO_KLRG1_LOW_EFFECTOR_CD8_TCELL_UP |
| GSE10325_CD4_ TCELL_VS_LUPUS_CD4_ TCELL_DN | CCR1\|SP100\|PTPN22\|MT1H\|ATP6V1A\|TRIM21\|PRPS2\|STAT1\|FBXO22\|CENPE\|EIF2AK2\|SELP\|TAPBP\|SNRPG\|SAMD9\|MX1\|OAS3\|RNASE2\|RALA\|PTPN2\|TFEC\|LAMP2\|LAMP3\|TNIP2\|CDC5L\|MT1X\|PCYT1A\|CAPN2\|UAP1L1\|TMSB10\|PLA2G7\|KPNA3\|HLA-DQA1\|LAP3\|PPIF\|IGKC\|IFI30\|CAPZA1\|FABP5\|PSME1\|CASP1\|ME2\|IFIH1\|ISG20\|NMI\|ISG15\|TRAFD1\|OASL\|GCH1\|PFKP\|MYD88\|DDX60\|IFIT1\|PLSCR1\|HNRNPD\|TRIM22\|TLR3\|UBE2L6\|TAP1\|MX2\|TBK1\|LGALS3BP\|PSMB9\|IFI16\|CASP7\|IFIT5\|PSMA6\|RBCK1\|GBP1\|KIAA0513\|IFI35\|OAS2\|RAB11FIP1\|COPS4\|PSME2 | 75 | 0.01 | GSE10325_CD4_TCELL_VS_LUPUS_CD4_TCELL_DN |

**Table.S2. sgRNA sequences.**

| **ID** | **Sequence** | **Annotation** |
| --- | --- | --- |
| SEQ ID NO:1 | CCCCAGCACTAACTGGTCAC | TargetSeq.1-ITPRIPL1-sgRNA1 |
| SEQ ID NO:2 | AGACATGGGGTGGCCGTTCC | TargetSeq.2-ITPRIPL1-sgRNA2 |
| SEQ ID NO:3 | TCCTGGCCATCGGCCTGGAA | TargetSeq.3-ITPRIPL1-sgRNA3 |
| SEQ ID NO:4 | CACCGCCCCAGCACTAACTGGTCAC | Primer-T1-sgRNA1 |
| SEQ ID NO:5 | AAACGTGACCAGTTAGTGCTGGGGC | Primer-B1-sgRNA1 |
| SEQ ID NO:6 | CACCGAGACATGGGGTGGCCGTTCC | Primer-T2-sgRNA2 |
| SEQ ID NO:7 | AAACGGAACGGCCACCCCATGTCTC | Primer-B2-sgRNA2 |
| SEQ ID NO:8 | CACCGTCCTGGCCATCGGCCTGGAA | Primer-T3-sgRNA3 |
| SEQ ID NO:9 | AAACTTCCAGGCCGATGGCCAGGAC | Primer-B3-sgRNA3 |

**Table.S3. Peptide and antibody sequences.**

| **ID** | **Sequence** | **Annotation** |
| --- | --- | --- |
| SEQ ID NO:1 | HPLMVSDRMDLDTLA | P1 |
| SEQ ID NO:2 | VSDRMDLDTLARSRQ | P2 |
| SEQ ID NO:3 | MDLDTLARSRQLEKR | P3 |
| SEQ ID NO:4 | TLARSRQLEKRMSEE | P4 |
| SEQ ID NO:5 | SRQLEKRMSEEMRLL | P5 |
| SEQ ID NO:6 | EKRMSEEMRLLEMEF | P6 |
| SEQ ID NO:7 | SEEMRLLEMEFEERK | P7 |
| SEQ ID NO:8 | RLLEMEFEERKRAAE | P8 |
| SEQ ID NO:9 | MEFEERKRAAEQRQK | P9 |
| SEQ ID NO:10 | ERKRAAEQRQKAENF | P10 |
| SEQ ID NO:11 | AAEQRQKAENFWTGD | P11 |
| SEQ ID NO:12 | RQKAENFWTGDTSSD | P12 |
| SEQ ID NO:13 | ENFWTGDTSSDQLVL | P13 |
| SEQ ID NO:14 | TGDTSSDQLVLGKKD | P14 |
| SEQ ID NO:15 | SSDQLVLGKKDMGWP | P15 |
| SEQ ID NO:16 | LVLGKKDMGWPFQAD | P16 |
| SEQ ID NO:17 | KKDMGWPFQADGQEG | P17 |
| SEQ ID NO:18 | MEWRIFLFILSGTAGVHSQVQLQQSGPELVKPGASVRMSCKTSGYTFTDYVISWVKQRPGQGLEWIGEIFPRTGSTYYNENFKATATLTADKSSNTAYMQLSSLTSEDSAAYFCAFITSVDWAMEYWGQGTSVTVSSAKTTPPSVYPLAPGSAAQTNSMVTLGCLVKGYFPEPVTVTWNSGSLSSGVHTFPAVLQSDLYTLSSSVTVPSSTWPSETVTCNVAHPASSTKVDKKIVPRDCGCKPCICTVPEVSSVFIFPPKPKDVLTITLTPKVTCVVVDISKDDPEVQFSWFVDDVEVHTAQTQPREEQFNSTFRSVSELPIMHQDWLNGKEFKCRVNSAAFPAPIEKTISKTKGRPKAPQVYTIPPPKEQMAKDKVSLTCMITDFFPEDITVEWQWNGQPAENYKNTQPIMDTDGSYFVYSKLNVQKSNWEAGNTFTCSVLHEGLHNHHTEKSLSHSPGK | 13B7A6H3 HC1 |
| SEQ ID NO:19 | MKLPVRLLVLMFWIPGSSSDVVMTQTPLSLPVSLGDQASISCRSSESLVNSKGNTHLHWYLQKPGQSPKLLIYKVSNRFSGVPDRFSGSGSGTDFTLKISRVEAEDLGVYFCSQSTHAPYTFGGGTKLEIKRADAAPTVSIFPPSSEQLTSGGASVVCFLNNFYPKDINVKWKIDGSERQNGVLNSWTDQDSKDSTYSMSSTLTLTKDEYERHNSYTCEATHKTSTSPIVKSFNRNEC | 13B7A6H3 LC1 |
| SEQ ID NO:20 | QVQLQQSGPELVKPGASVRMSCKTSGYTFTDYVISWVKQRPGQGLEWIGEIFPRTGSTYYNENFKATATLTADKSSNTAYMQLSSLTSEDSAAYFCAFITSVDWAMEYWGQGTSVTVSS | 13B7A6H3 VH1 |
| SEQ ID NO:21 | DVVMTQTPLSLPVSLGDQASISCRSSESLVNSKGNTHLHWYLQKPGQSPKLLIYKVSNRFSGVPDRFSGSGSGTDFTLKISRVEAEDLGVYFCSQSTHAPYTFGGGTKLEIKR | 13B7A6H3 VL1 |
| SEQ ID NO:22 | GYTFTDYV | 13B7A6H3 HCDR1 |
| SEQ ID NO:23 | IFPRTGST | 13B7A6H3 HCDR2 |
| SEQ ID NO:24 | AFITSVDWAMEY | 13B7A6H3 HCDR3 |
| SEQ ID NO:25 | ESLVNSKGNTH | 13B7A6H3 LCDR1 |
| SEQ ID NO:26 | KV | 13B7A6H3 LCDR2 |
| SEQ ID NO:27 | SQSTHAPYT | 13B7A6H3 LCDR3 |

**Supplementary Figures**

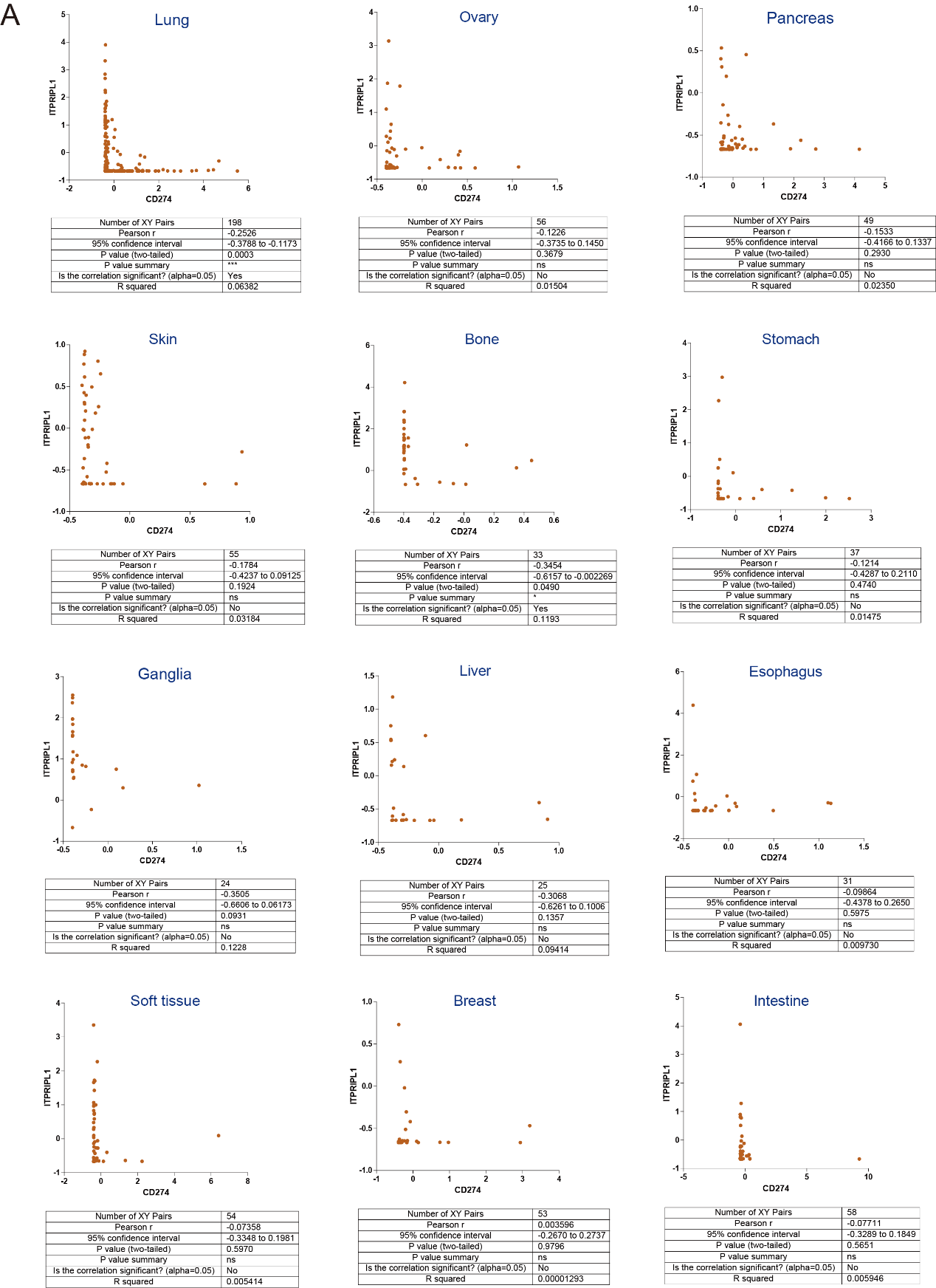

**Fig.S1.** A, CCLE analysis showing mutually exclusive expression pattern between ITPRIPL1 and CD274 in different cancers.

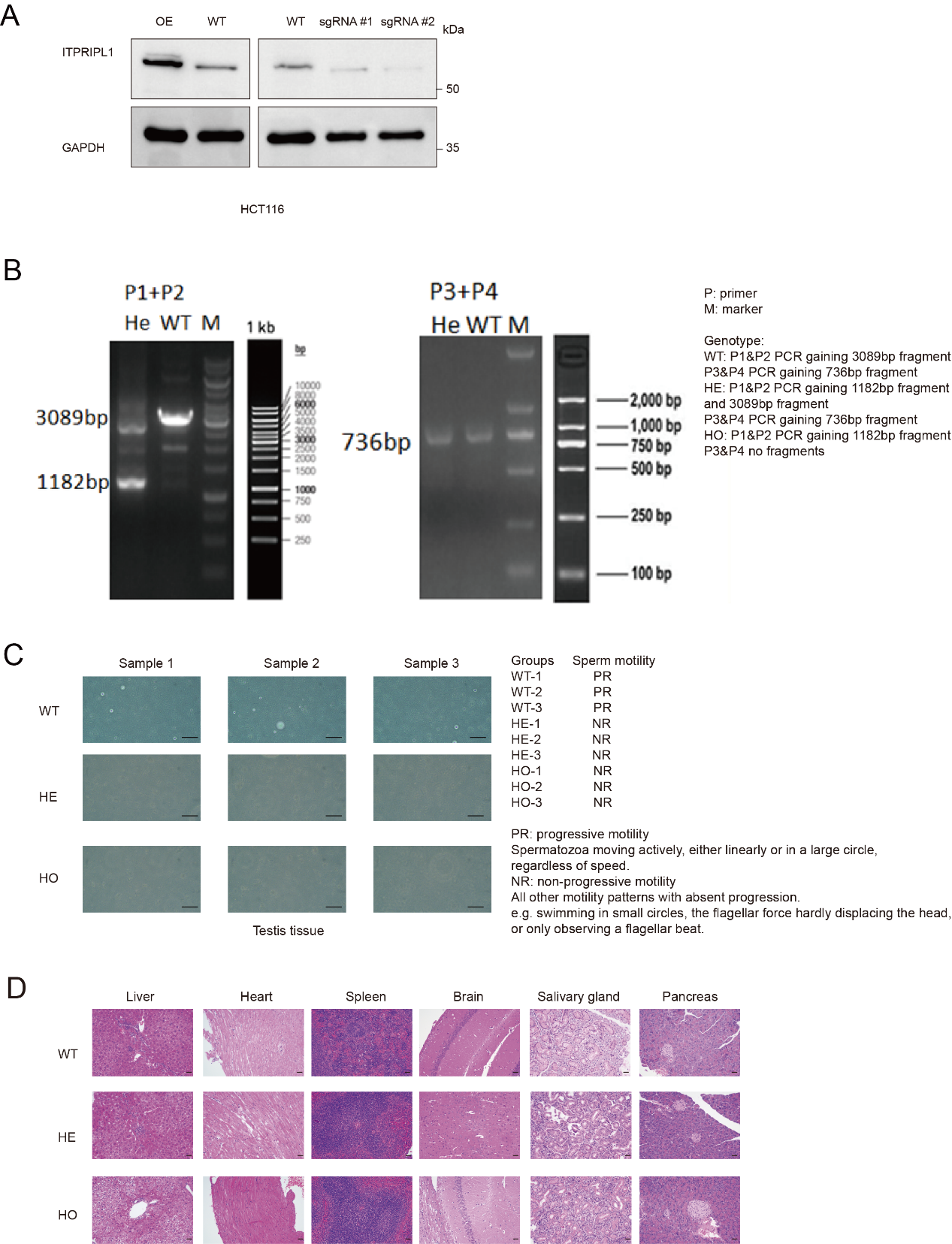

**Fig.S2.** A, Immunoblot showing the establishment of HCT116-ITPRIPL1 OE/KO cells (n=3). B, PCR results showing the ITPRIPL1 knockout status in ITPRIPL1 HO mice vs WT mice. C, Microscopic views (100X) of sperm from different groups of mice and dynamic evaluation according to WHO clinical guidance (n=3). D, Microscopic views (100X) of HE stains of different organ tissues apart from testis showing no obvious abnormalities in ITPRIPL1 HE/HO groups.

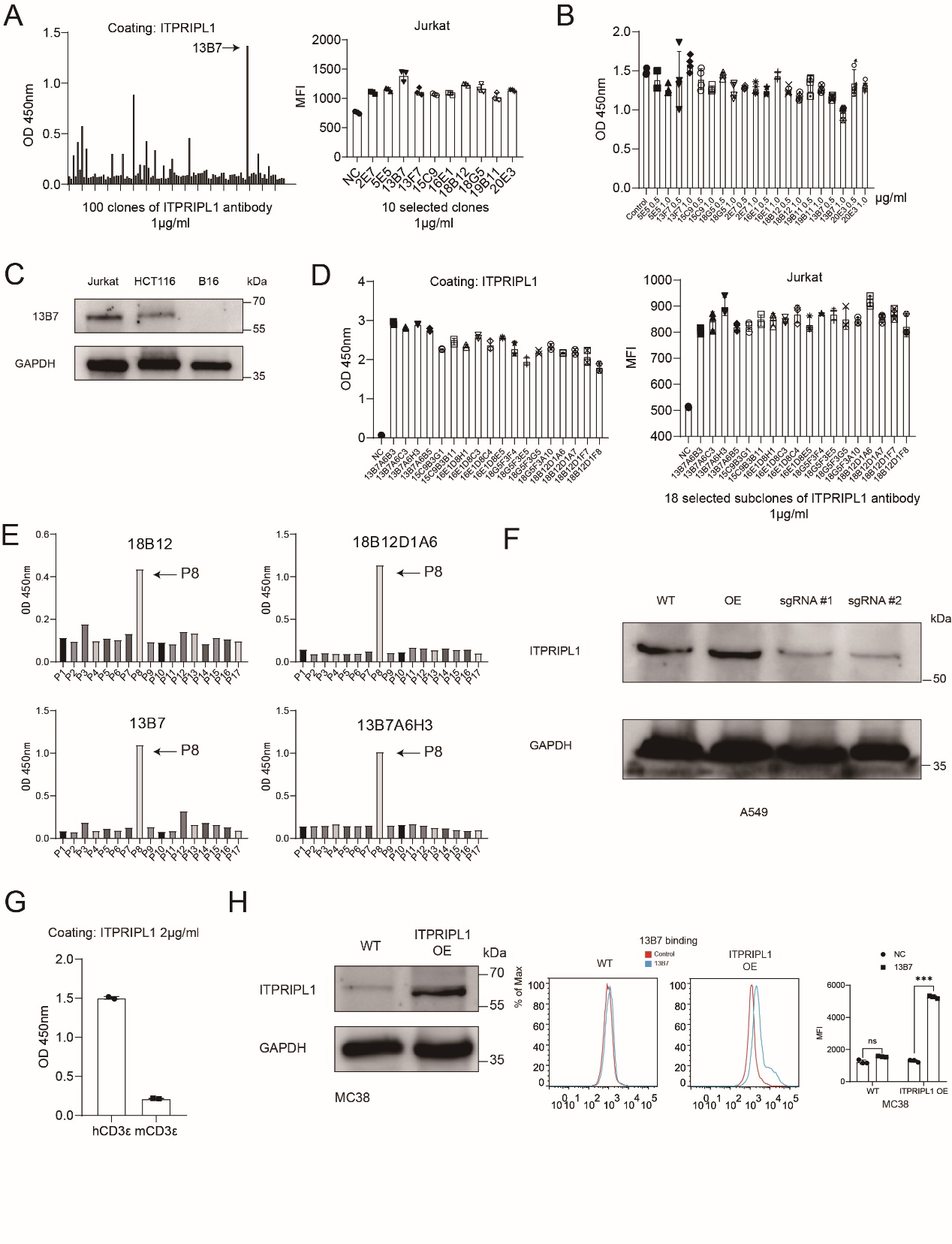

**Fig.S3.** A, ELISA showing the best affinity of 13B7 among 100 antibodies; FACS showing the best affinity of 13B7 among 10 selected clones (n=3). B, ELISA showing the ability of 10 selected clones to block ITPRIPL1-CD3ε interaction, with 13B7 showing the greatest effect (n=4). C, Western blot results by 13B7 antibody, showing a relatively specific band at the approximate molecular weight position of ITPRIPL1. D, ELISA (n=2) and FACS (n=3) showing the binding affinity of 18 selected subclones. E, ELISA showing the specific peptide region P8 as the binding part of 13B7 and 18B12 clones. F, Immunoblot showing the establishment of ITPRIPL1 OE and KO A549 cell groups (n=3). G, ELISA showing ITPRIPL1 interacted with human CD3ε but not mouse CD3ε (n=2). H, Immunoblot and FACS showing the establishment of MC38-ITPRIPL1 OE cell lines (n=3). Data are mean ± s.d. **P<0.05, ***P<0.001, ****P<0.0001. Two-tailed Student’s t-test.
